# Ultra-long-range Polycomb-coupled interactions underlie subtype identity of human cortical neurons

**DOI:** 10.1101/2025.11.05.686502

**Authors:** Ilya A. Pletenev, Nikita Vaulin, Maria N. Molodova, Ekaterina Kuznechenkova, Anastasia Soldatenkova, Olga I. Efimova, Anna V. Tvorogova, Philipp Khaitovich, Sergey V. Razin, Sergey V. Ulianov, Ekaterina E. Khrameeva

**Author notes:** These authors contributed equally to this work (Ilya A. Pletenev, Nikita Vaulin).

## Abstract

Regulation of gene expression by Polycomb group (PcG) proteins orchestrates neural development and cell-type specification. However, the mechanisms by which neural progenitor cells diversify into the myriad cell types of the human brain remain poorly understood. Here, we investigate the role of Polycomb proteins in shaping the single-cell 3D genome, transcriptome, and chromatin binding landscape of the developing and adult human cortex. We show that PcG proteins establish a network of long-range repressive interactions encompassing loci with neuronal marker genes. These interactions occur in a neuron-type–dependent manner, selectively targeting genes that must remain silenced in a given cell type. These interactions are present in fetal neurons and increase markedly in strength after birth. Our results highlight PcG proteins as key regulators of neuronal cell fate specification in the human brain.

## 1 Introduction

During organism development, cells undergo differentiation which is orchestrated by transcription factors. Key players of developmental regulation are Polycomb-group (PcG) proteins — a set of transcription factors, histone-modifying enzymes, and associated cofactors that assemble into two Polycomb repressive complexes (PRC1 and PRC2) whose principal function is transcriptional repression [Blackledge and Klose, 2021, Chittock et al., 2017]. Both complexes possess “reader” and “writer” activities: PRC1 deposits H2AK119ub1, while PRC2 deposits H3K27me3, and they can bind both these modifications [Blackledge and Klose, 2021]. Moreover, PcG proteins mediate the formation of higher-order 3D chromatin structure that enables Polycomb spreading [Kraft et al., 2022, Oksuz et al., 2018] and ensure long-term maintenance of repressive states [Owen et al., 2023, Kim et al., 2023].

PcG proteins are critical regulators of cortical development, including dorsoventral patterning of the telencephalon [Eto et al., 2020], cortical neurogenesis [Hirabayashi et al., 2009, Pereira et al., 2010, Tsuboi et al., 2018], and neuronal subtype specification [Morimoto-Suzki et al., 2014]. However, the contribution of Polycomb-mediated 3D chromatin interactions to these processes remains poorly understood. Most studies have focused on *in vitro*-differentiated neurons or neuronal progenitor cells [Bonev et al., 2017, Kundu et al., 2017, Cruz-Molina et al., 2017, Crispatzu et al., 2021], with few investigations addressing the 3D organization of H3K27me3-occupied loci in the adult cortex [Takei et al., 2021, Lu et al., 2022].

We and others have shown that the three-dimensional (3D) genome of human neurons is distinct from that of other brain and non-brain cell types, both in bulk and at single-cell resolution [Rahman et al., 2023, Hu et al., 2021, Pletenev et al., 2024, Tian et al., 2023, Heffel et al., 2024, Zhou et al., 2025]. Specifically, in mature neurons, many genes encoding transcription factors involved in neuronal differentiation — including *SOX2* and *DLX2* — are marked by H3K27me3 [Pletenev et al., 2024, Von Schimmelmann et al., 2016, Sawai et al., 2022]. Moreover, unlike other brain cell types, these loci in neurons engage in exceptionally long-range chromatin interactions (hereafter referred to as PcG contacts), spanning up to several hundred megabases [Pletenev et al., 2024, Vieux-Rochas et al., 2015]. The functional significance of these contacts, however, remains unclear.

To investigate how Polycomb-associated 3D chromatin architecture influences neuronal development and function, we analyzed publicly available single-cell datasets profiling chromatin structure in neurons from the post-mortem developing and adult human cortex. We show that PcG-associated contacts are organized in a neuron type-and developmental stage-specific manner. To map genome-wide binding of RING1B, the core subunit of PRC1, we performed ChIP-seq in human neurons. We find that core PRC1 subunits are highly enriched at a small subset of gene promoters that engage in long-range interactions with one another.

## 2 Results

### 2.1 Many PcG contacts are neuron-type-specific

The 3D genome varies significantly across neuronal subtypes of the human cortex at the level of contact distance distribution, chromatin compartments, topologically associating domains, and chromatin loops [Tian et al., 2023, Heffel et al., 2024]. Previous studies have analyzed chromatin loops spanning less than 10 Mb, which are predominantly CTCF-mediated. As we have previously demonstrated [Pletenev et al., 2024], PcG contacts differ markedly from CTCF-mediated loops: their anchors overlap H3K27me3 domains, connect regions at greater genomic distances, and form both intra-and interchromosomal interactions (Fig. 1A). Therefore, we aimed to investigate the cell-type specificity of PcG contacts using publicly available 3D chromatin data from the human cerebral cortex [Tian et al., 2023]. We first examined excitatory (EN) and inhibitory (IN) neurons, the two major classes that show the most pronounced chromatin differences in the cortex [Tian et al., 2023]. We performed manual and automated annotation of loci that form PcG contacts (“PcG anchors”) using EN and IN contact maps and selected the union of the anchors found (Methods, Fig. S1A). This procedure yielded 262 PcG anchors encompassing many neuronal transcription-factor and effector genes (Table S1). From all intrachromosomal pairwise interactions between these loci, we retained those with intensity greater than background (Methods). We then calculated normalized PcG-contact intensity in EN, IN, and non-neurons (NN) from snm3C–seq data (Methods). K-means clustering reveals that most contacts have high signal in both EN and IN, but display a preference for one of the two cell types (Figs. 1B, C; clusters “EN-prevalent” and “IN-prevalent”). By contrast, 196 contacts are specific to either EN or IN (Figs. 1B, C; clusters “EN-specific” and “IN-specific”), consistent with differential H3K27me3 ChIP–seq signal at the corresponding loci (Fig. S1B). As stated above, PcG contacts are different from CTCF-mediated loops (Fig. S1C). For some PcG anchors, CTCF signal is observed at the boundaries of the region, which agrees with reported findings that CTCF restricts Polycomb domain spreading [Weth et al., 2014].

**Fig. 1.**
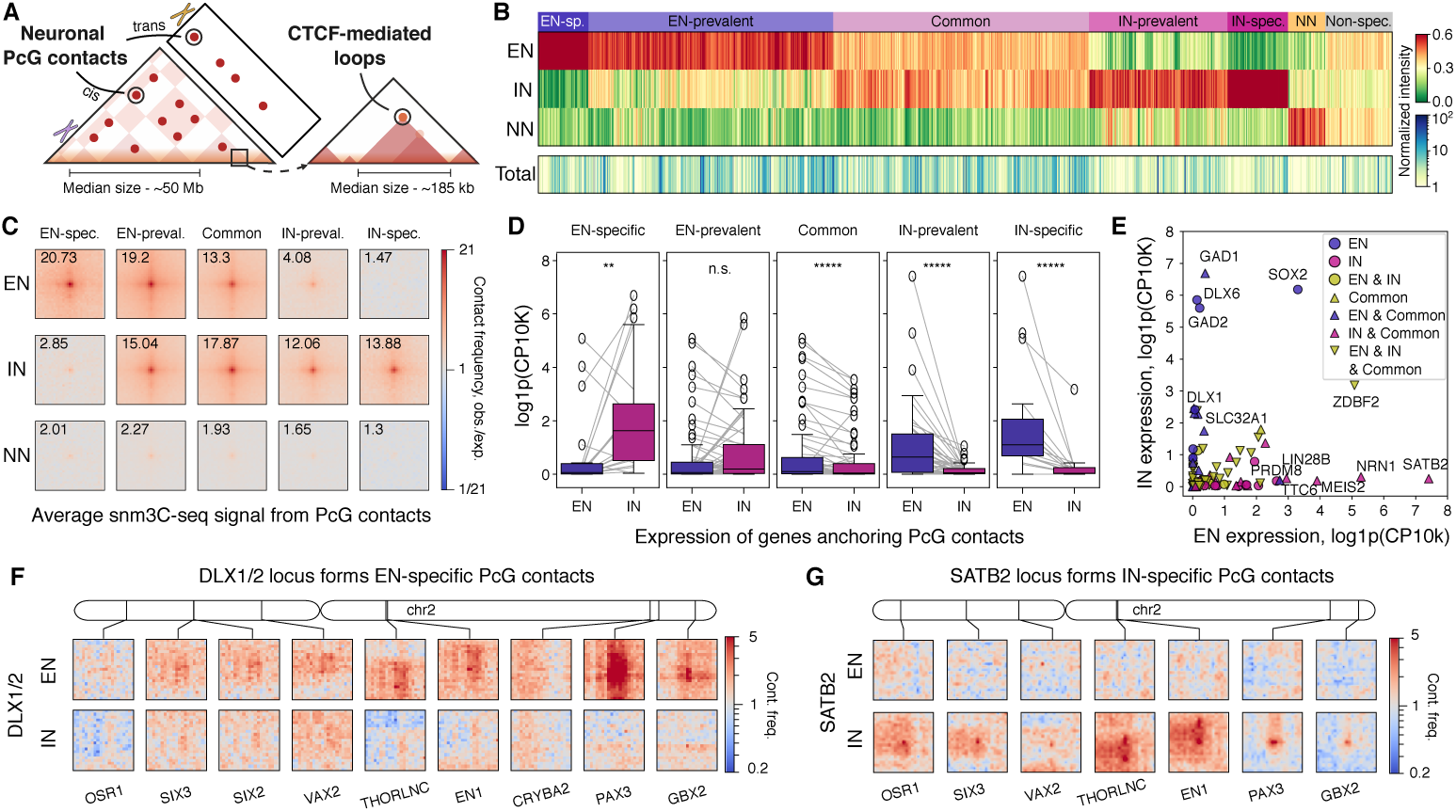
A. Schematic representation of the difference between neuronal PcG contacts and CTCF-mediated loops. Median size for PcG loops is taken from [Pletenev et al., 2024], for CTCF-mediated loops — from [Rao et al., 2014]. B. Normalized PcG contact frequency calculated in single cells and aggregated into excitatory (EN), inhibitory neurons (IN), and non-neurons (see Methods). Each column corresponds to a separate PcG contact and is normalized to one.”Total” row corresponds to PcG contact frequency without normalization and represents the average intensity of a PcG contact C. Average PcG contact frequency per cluster calculated from pseudobulk snm3C-seq data and normalized by expected signal at a given genomic distance. D. Expression of genes that overlap PcG anchors of contacts from specific clusters. ** P-value ¡ 0.01, ***** P-value ¡ 0.00001, two-sided Wilcoxon rank-sum test. E. Same data as in D; each contact corresponds to a gene, color represents clusters in which gene overlaps PcG anchor. “EN” — “EN-specific” or “EN-prevalent”, “IN” — “IN-specific” or “IN-prevalent”. We removed 7 genes and one transcript from the plot: *WDR41*, *RSRC1*, *CCNC*, *OSTM1*, *NUP133*, *ETFA*, *STK3*, and *AC058822.1* — under closer examination, these genes were found close to, but not overlapping PcG anchor boundaries. F,G. Snippets of aggregated snm3C-seq contact matrices showing differential PcG contacts between EN and IN for SATB2 (F) and DLX1/2 (G) loci. A gene name at the bottom of each plot indicates the gene closest to the contact anchor; only genes silent in both cell types were considered.

A single genomic region can serve as an anchor for multiple PcG contacts (Fig. S1D). We next asked whether all contacts formed by a given anchor belong to the same PcG contact cluster. We found that “EN-prevalent” and “IN-prevalent” contacts tend to share a common set of anchors, whereas “EN-specific” and “IN-specific” clusters each utilize distinct anchors (Fig. S1E). Clusters appearing to contain “NN-specific” or “Non-specific” PcG contacts, upon closer inspection, exhibited low contact matrix values (Fig. 1B, “Total” track; Fig. S1F) and were therefore considered false positives.

We have previously described cases where PcG contact intensity anticorrelates with the expression of the anchoring gene [Pletenev et al., 2024]. However, a systematic analysis of this relationship is complicated by the fact that a single PcG contact brings together two broad genomic loci, each typically containing multiple genes. To address this, we leveraged publicly available snRNA-seq data from the adult human brain [Siletti et al., 2023]. Within each cluster, we selected PcG contacts that bring together one anchor containing at least one gene expressed in neurons (“selectively expressed” locus) and another anchor with no expression. We refer to such interactions as “representative”. At these contacts, the expression of active genes anticorrelates with contact intensity (Figs. 1D, E).

We next examined differentially expressed (DE) genes within each cluster. The “EN-specific” cluster includes previously reported [Pletenev et al., 2024] PcG contacts with genes encoding GABAergic transcription factors — *DLX5*, *DLX6*, and *SOX2* — as well as functional markers such as *GAD2* and *SLC32A1*. It also contains newly identified contacts with transcription factors (*DLX1*, *DLX2*, *SP5*, *SP9*, *LHX6*) and the functional marker *GAD1*. Several genes anchored by “EN-specific” PcG contacts participate in shared regulatory pathways governing the development and specification of GABAergic neurons. For example, in postmitotic neurons, DLX1 and DLX2 induce the expression of *GAD1*, *DLX5*, *DLX6*, and *SP9*, while *SP9* expression is required for expression of *LHX6* during the migration and maturation of GABAergic neurons [Lim et al., 2018]. Representative “EN-specific” contacts are illustrated by *DLX1/2* - mediated contacts in Figure 1F.

Symmetrically, the “IN-specific” cluster contains transcription factor genes — *SATB2*, *TBR1*, and *NR4A2* — related to glutamatergic neuron specification. For example, *SATB2* and *TBR1* mutually regulate each other to establish neuronal identity in concert with *FEZF2*, which is classified within the “common” cluster [Srinivasan et al., 2012, McKenna et al., 2015]. Representative “IN-specific” contacts are illustrated by *SATB2*-mediated contacts in Figure 1G.

These results indicate a potential role of PcG contacts in the specification of human cortical neurons.

### 2.2 Some PcG contacts distinguish specific neuronal subtypes

Excitatory and inhibitory neurons in the cerebral cortex are heterogeneous and can be subdivided into distinct subtypes based on features such as location, developmental origin, and projection targets [Mao and Staiger, 2024]. These distinctions are reflected in their transcriptomes [Siletti et al., 2023], with many transcription factors showing subtype-specific expression patterns [Yao et al., 2023]. To extend our analysis, we quantified PcG contact occurrence across neuronal subtypes and found strong subtype specificity (Methods, Fig. S2A).

Previous work has shown that human brain cell types can be distinguished by 3D-genome features such as chromatin compartments, TADs, and short-range (¡5 Mb) loops [Tian et al., 2023]. We next asked whether brain cell types could also be distinguished by PcG contacts. This was not feasible at the single-cell level due to the low number of reads corresponding to PcG contacts, so we aggregated signal by cell type. Principal component analysis (PCA) reveals the separation of cell types into neurons and non-neurons (PC1) and a further separation of neurons into EN and IN (PC2; Fig. 2A). PC1 reflects average intensity of PcG contacts, as PC1 loadings are mostly positive (Fig. S2B). Interestingly, neuronal subtypes do not cluster together but instead spread along PC1. Dot positions are robust across biological samples (Fig. S2C), suggesting that subtype-specific factors contribute to average PcG contact intensity.

**Fig. 2.**
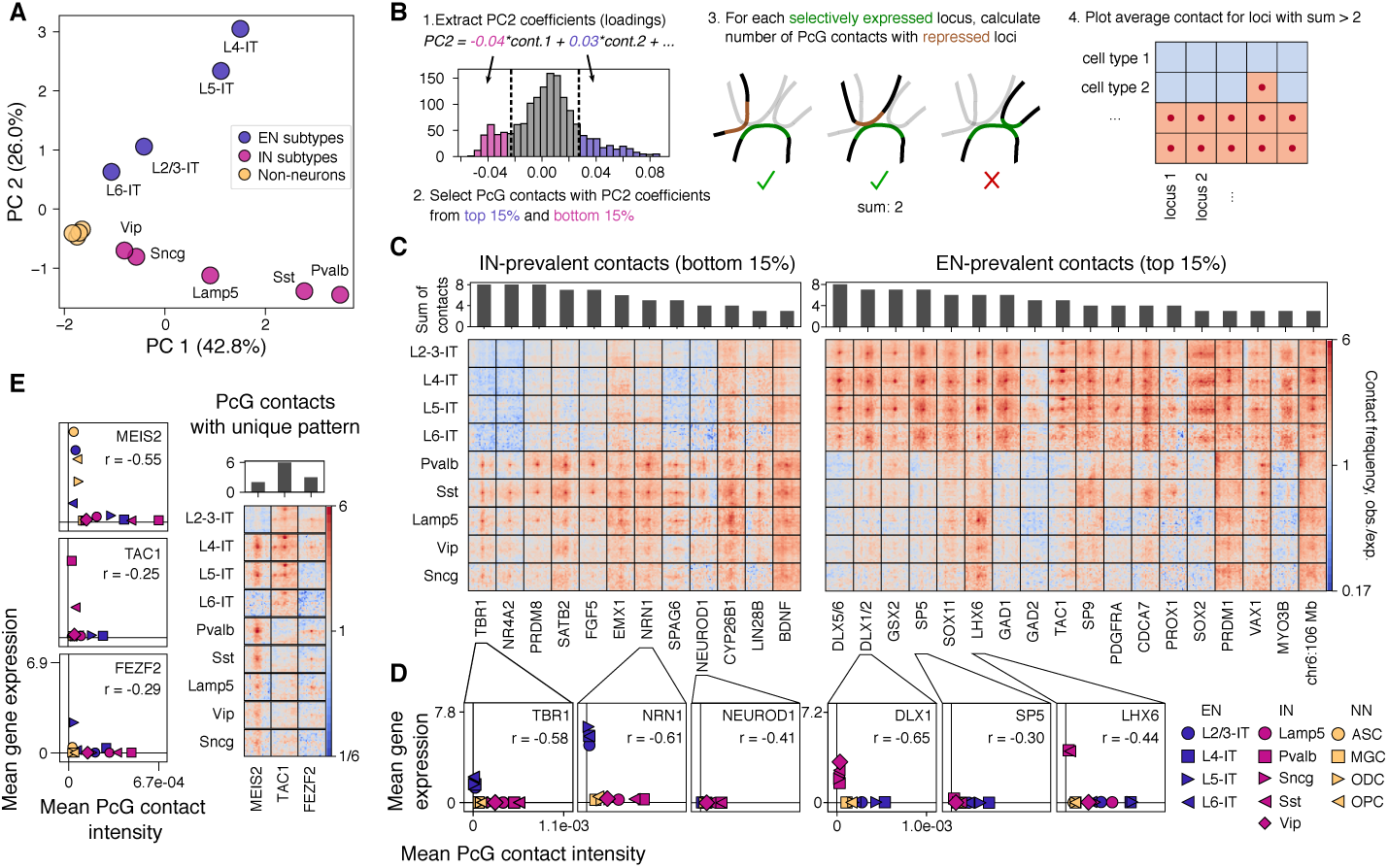
A. Principal component analysis based on PcG contact intensity calculated for neuronal subtypes and non-neurons. Color depicts type of cells: EN, IN, or non-neurons grouped into oligodendrocytes, oligodendrocyte precursor cells, astrocytes, and microglia. B. Steps of the analysis of EN-prevalent and IN-prevalent PcG contacts shown on (C). C. Averaged EN-prevalent and IN-prevalent PcG contacts formed by a given anchor. Contact matrix resolution — 100 kb. Region length — 2 Mb. Signal is normalized by expected at a given distance. Color scale is identical for all contacts. Grey bars on top represent the number of PcG contacts used for averaging. D. Examples of how mean PcG contact intensity relates to mean gene expression of a gene overlapping PcG anchor. Pearson correlation coefficient (r) is shown. “ASC” — astrocytes, “MGC” — microglia, “ODC” — oligodendrocytes, “OPC” — oligodendrocyte progenitor cells. E. Examples of PcG anchors with unique neuronal subtype specificity. Grey bars on top represent the number of PcG contacts used for averaging. Scatter plots below show examples of how mean PcG contact intensity relates to mean gene expression of a gene overlapping PcG anchor. Dot colors and shapes are same as on (D).

We next asked which PcG anchors contributed most to the separation of EN and IN in the PCA. To identify these anchors, we selected PcG contacts with the highest contribution to PC2 variance and retained “representative” ones (Fig. 2B). We then retained anchors that formed at least three such contacts to ensure robustness and averaged the resulting contacts (Fig. 2C). As in Figure 1D, expression of genes overlapping PcG anchors (“anchor genes”) anticorrelates with contact intensity across neuronal subtypes (Fig. 2D, Fig. S2D). Moreover, most genes in Figure 2D demonstrate a mutually exclusive relationship between gene expression and the presence of PcG contact.

Although most contacts in Figure 2C are EN-or IN-specific, more intricate contact patterns are also observed. For example, the *LHX6* anchor forms contacts in EN and in three out of five IN subtypes, but not in Pvalb or Sst, consistent with the known activity of *LHX6* in these subtypes [Lim et al., 2018]. To explore such patterns, we analyzed the relationship between expression and contact intensity for all PcG anchors and selected differentially expressed anchors in which points were not grouped simply into EN and IN (Methods). Three examples are shown in Fig. 2E, each consistent with known roles of the encoded proteins. The transcription factor FEZF2 controls deep-layer neuron specification [Heavner et al., 2020], while the neurotransmitter precursor TAC1 is a marker of Pvalb neurons [Pfeffer et al., 2013]. The transcription factor MEIS2 is involved in the specification of striatal medium spiny neurons [Su et al., 2022], dopaminergic neurons [Agoston et al., 2014], and serotonin receptor 3A interneurons [Frazer et al., 2017]; however, to our knowledge, its role in glutamatergic cortical neurons has not been previously described.

### 2.3 PcG contacts emerge during brain development

Many TF genes that are inactive and anchored by PcG contacts in adult brain are expressed during development. For example, the expression of *BHLHE22* decreases monotonically in cortical neurons from mid-gestation to adulthood [Herring et al., 2022] suggesting that PcG-contact intensity might also vary across stages. To investigate this, we reanalyzed publicly available snm3C-seq data from the developing human nervous system at four time points: second trimester (2T), third trimester (3T), infant, and adult cerebral cortex [Heffel et al., 2024], using cell-type annotations from the original paper.

First, we analyzed the 3D genome of radial glial cells (RG) — progenitors that form neurons and glia during mid-gestation [Leibovitz et al., 2022]. It was recently shown that RG separate into two clusters based on their 3D chromatin structure: one closer to neurons (“neuronal”) and the other to astrocytes (“glial”) [Heffel et al., 2024]. We find that PcG contacts are already present in “neuronal” RG but absent in “glial” RG (Fig. S3A), indicating that a bright PcG-contact pattern is a distinguishing feature of neurons that appears during RG differentiation in the second trimester of human development.

We continued by studying the intensity of PcG contacts during the development of neurons. Contact frequency was first measured for genomic regions annotated as PcG contacts in the adult brain, revealing that PcG-contact intensity increases substantially after birth in both EN and IN (Figs. 3A, B, Fig. S3B). In the fetal brain, intensity changes in a cell-type dependent manner; in particular, PcG contacts at 2T are much more pronounced in EN than in IN (Fig. 3B, Fig. S3C). PCA using contact intensities as features demonstrates clear separation of samples by age (PC1) and cell type (PC2) (Fig. 3C), with most features (83%) showing positive PC1 loadings, reflecting an age-related increase in average contact intensity (Fig. S3D). PC2 loadings span both positive and negative values, indicating the existence of EN-specific and IN-specific PcG contacts not only in adult but also in fetal and infant brains, as exemplified in Figure 3D.

**Fig. 3.**
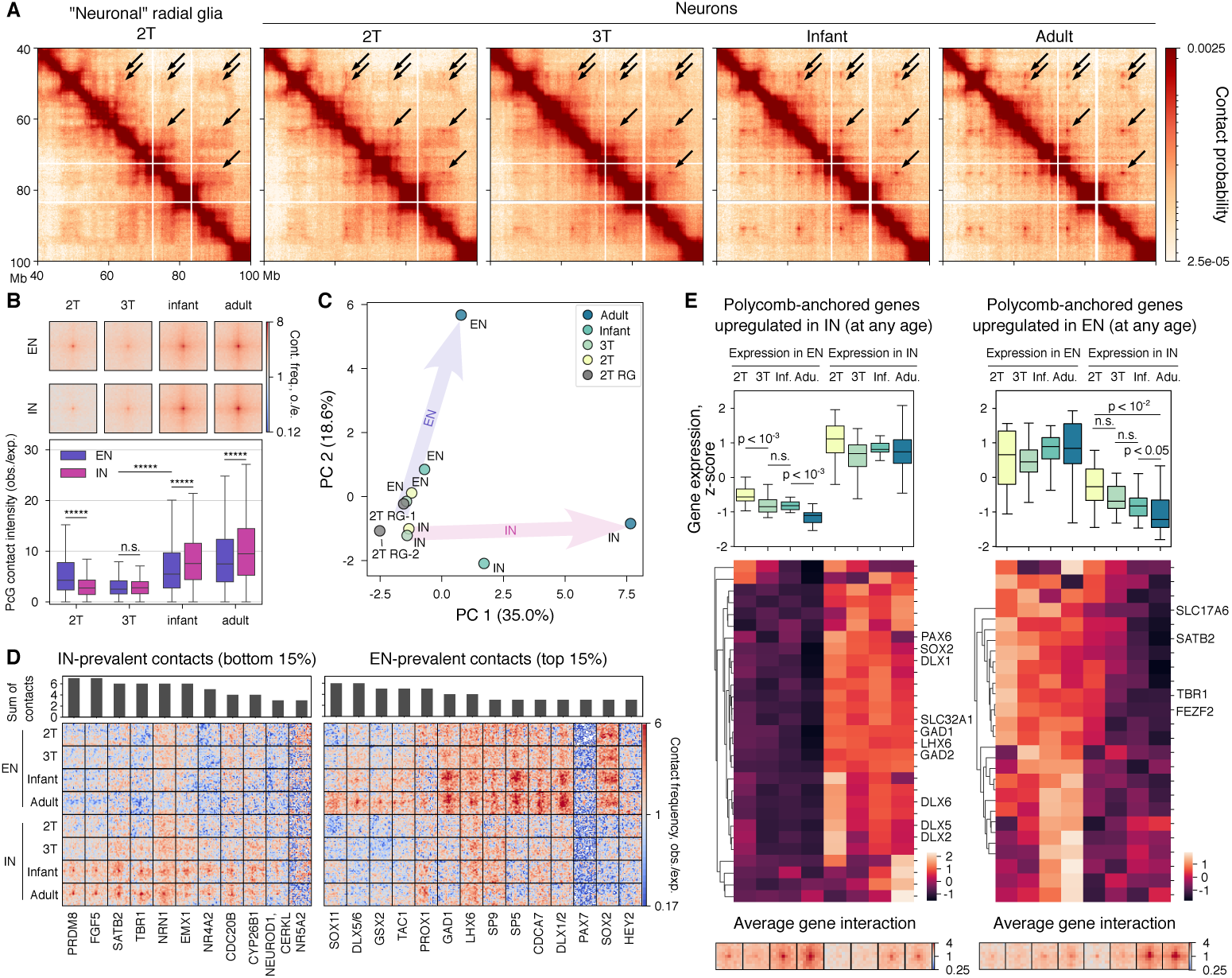
PcG contacts emerge during neuronal maturation. A. Example of a pseudobulk snm3C-seq matrix region with PcG contacts that gradually appear during neuronal maturation. B. The average snm3C-seq (pseudobulk) signal (observed over expected) of PcG contacts for neurons grouped into EN and IN (top) and corresponding signal distribution of the central pixel (bottom). ***** P-value ¡ 0.00001, two-sided Wilcoxon test. C. Principal component analysis based on PcG contact intensity calculated for neuronal subtypes and different stages of development. Color depicts stage of development. D. Averaged EN-prevalent and IN-prevalent PcG contacts formed by a given anchor. Contact matrix resolution — 100 kb. Region length — 2 Mb. Signal is normalized by expected at a given distance. Color scale is identical for all contacts. Grey bars on top represent the number of PcG contacts used for averaging. Analysis was performed similarly to Figs. 2C. E. Transcription level of PcG anchor genes differentially expressed between EN and IN at any stage of development. (bottom) Average PcG contact for a given set of differentially expressed genes.

Because activating an anchoring gene can eliminate its PcG contact, we hypothesized that stage-dependent changes in average intensity reflect differential expression of anchor genes. To examine this, we utilized publicly available snRNA-seq data on human cortical development [Herring et al., 2022], organizing gene expression data by four time points to align with snm3C-seq. We find that 44% of anchoring genes (148 out of 264) are expressed in at least one of the considered stages (Fig. S3E). This suggests that PcG contacts may contribute to transcriptional repression during neuronal maturation. Notably, even contacts anchored by genes that remain consistently repressed throughout development changed in intensity in a similar way to all PcG contacts (Fig. S3F), implying a genome-wide mechanism rather than locus-specific regulation.

Figure 3D demonstrates that some EN-specific and IN-specific PcG contacts observed in the adult brain emerge earlier during development and increase in intensity with age. These contacts anchor many markers of EN and IN differentiation, such as *DLX1/2* and *SATB2*. Although each marker is expressed in its corresponding neuronal type, a progressive increase in PcG-contact intensity in the other cell type may indicate a gradual decline in gene expression as cell-type identity is established. To test this hypothesis, we analyzed the expression of marker genes anchored by PcG contacts. Candidate genes were identified by differential expression analysis between EN and IN at each developmental stage, yielding two non-overlapping sets of PcG-anchored genes: those upregulated in EN at one or more stages and those upregulated in IN at one or more stages. Interestingly, whereas many IN-specific genes (23/33) become differentially expressed during fetal development, most EN-specific genes (21/28) acquire differential expression only after birth (Fig. S3G). Comparing expression across stages revealed that genes anchored by a PcG contact in a certain cell type undergo gradual repression with age in that cell type (Fig. 3E). Together, these findings suggest that increasing PcG-contact intensity may contribute to the progressive establishment of neuronal subtype identity.

We next asked whether maturing neurons form unique PcG contacts that are absent in adult neurons. Visual analysis of aggregated (pseudobulk) snm3C-seq maps did not reveal additional PcG contacts at 2T or 3T. To investigate further, we used publicly available fetal H3K27me3 ChIP-seq data from cortical plate at 2T [Rahman et al., 2023]. We calculated contact intensity between all intrachromosomal pairs of fetal-specific H3K27me3 regions separated by at least 3 Mb, and compared the resulting distribution of contact frequencies with PcG contacts annotated in adult neurons. We observe that fetal-specific H3K27me3 regions form, on average, significantly less frequent interactions than PcG contacts annotated in adult stage (Fig. S4A).

Differential PcG-contact intensity may reflect changes in PRC1 and PRC2 composition during development, as reported for earlier developmental stages [Liu et al., 2023]. Indeed, several Polycomb subunits show stage-dependent expression (Fig. S4B). To better understand these differences, we grouped Polycomb subunits by interchangeable paralogs that can alter properties of the complex [Tamburri et al., 2024, Brown et al., 2023, Niekamp et al., 2024] (Fig. S4C). We notice drastic changes in relative expression in some groups of paralogs, e.g., decreasing *EZH2* relative to *EZH1*. In the CBX group, *CBX2*, *CBX4* and *CBX8*, expressed at 2T, are gradually replaced by *CBX6* and *CBX7* in adult neurons, and *CBX2* and *CBX8* themselves are repressed by PcG contacts in adult neurons. Altogether, these results demonstrate that intensity of PcG contacts is altered during human cortical development and that genomewide chromatin reorganization, anchor gene activation, and differential expression of Polycomb genes might contribute to this phenomenon.

### 2.4 PcG contacts are characterized by PRC1 binding

We previously demonstrated that neuronal long-range contacts overlap H3K27me3 ChIP-seq signal, a histone mark deposited by Polycomb Repressive Complex 2 (PRC2) [Pletenev et al., 2024]. However, PRC1 is more directly associated with long-range chromatin interactions due to its ability to polymerize and form liquid condensates [Boyle et al., 2020, Eskeland et al., 2010, Tatavosian et al., 2019, Kundu et al., 2017, Blackledge and Klose, 2021], and PRC1 binding sites do not always overlap with H3K27me3 domains [Loubiere et al., 2020]. To gain more insight into the positioning of Polycomb complexes at PcG anchors, we performed ChIP-seq for RING1B on nuclei isolated from post-mortem human cortex and sorted by the neuronal marker NeuN. In addition, we leveraged publicly available ChIP-seq data for H3K27me3 and the PRC1 subunit BMI1 in adult neurons and oligodendrocytes [Loupe et al., 2024].

Since most of the PcG anchors overlap gene transcription start sites (TSSs), as demonstrated previously [Pletenev et al., 2024], we selected all TSSs showing a signal peak for at least one antibody in any cell type. We then calculated total ChIP-seq signal in a 40 kb window around each TSS and performed k-means clustering on these values (Methods). The resulting clusters reveal surprisingly different patterns of ChIP-seq signal (Fig. S5). Clusters 1 and 2 exhibit the highest and broadest enrichment for H3K27me3, BMI1, and RING1B, spanning up to 40kb, and show the highest overlap with PcG anchors (Fig. S5, left annotation track). The corresponding loci overlap long CpG islands and contain transcriptionally repressed genes (Fig. S5). The signal in neurons is significantly higher than in non-neuronal control, either oligodendrocytes or NeuN(-) cells (Fig. 4A). Thus, PcG contacts are characterized by the broad regions occupied by PRC1 and H3K27me3.

**Fig. 4.**
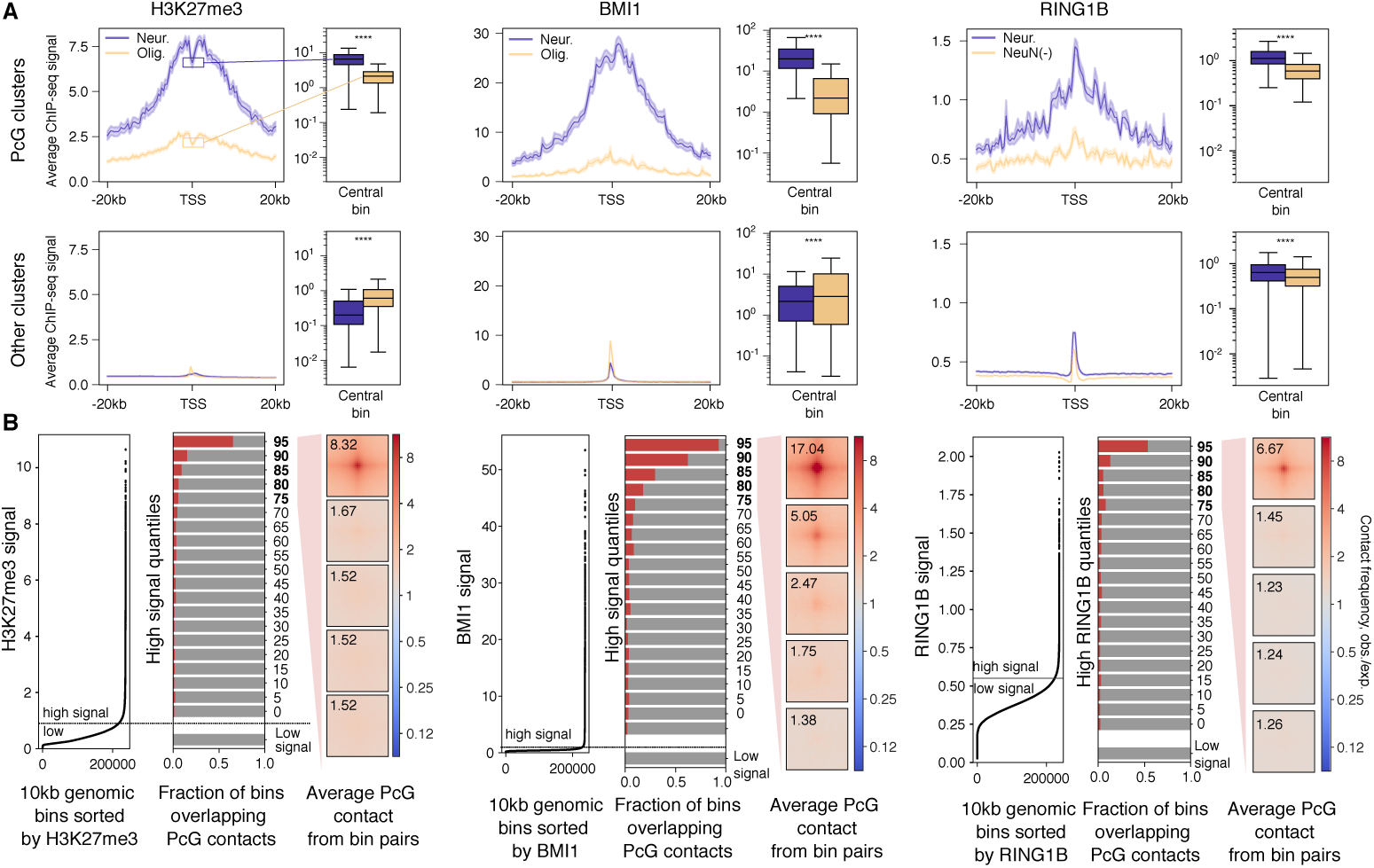
PcG contacts correspond to regions of high and broad ChIP-seq signal. A. Average ChIP-seq signal (-log10(Poisson P-value)) for combined PcG clusters. Area around the line represents standard error of the mean. Boxplots represent ChIP-seq signal at the central bin. **** P-value ¡ 0.0001, two-sided Wilcoxon test. B. Annotation-independent analysis of PcG contacts for H3K27me3 (left), BMI1 (center), and RING1B (right). (Left) A genome-wide distribution of ChIP-seq signal split in 10 kb genomic bins. The signal is grouped into “high” and “low” categories based on a threshold. (Center) Fraction of genomic bins overlapping PcG contact annotation calculated for “high” signal genomic bins split into 20 quantiles. Additionally, the fraction is calculated for all “low signal” bins. (Right) Average contact frequency calculated for the top 5 quantiles with “high signal” regardless of PcG contact annotation. Region size — 300 kb. Value in the corner represents mean value within a central 5×5 pixel area.

To assess the genome-wide distribution of PRC1 and H3K27me3 in neurons without relying on prior annotations, the genome was split into 10 kb bins and bins were ordered by ChIP–seq signal for each antibody (Fig. 4B, left). Distributions for all three antibodies showed prominent inflection points that we used to group genomic bins into”low signal” and”high signal”. We further focused on”high signal” bins and calculated the overlap with PcG anchors (Fig. 4B, center). Among “high-signal” bins the highest overlap with PcG anchors occurred in the top quantiles for H3K27me3, BMI1, and RING1B. Next, for each antibody we selected top 20% of”high signal” bins and computed average contact frequency between their pairs (regardless of PcG-contact annotation) (Fig. 4B, right). Average contact signal is the highest in the highest-signal groups, providing evidence that the very high PRC1 and H3K27me3 signal is associated with long-range PcG contact formation.

We then investigated what defines the high and broad binding of PRC1 and PRC2 at PcG anchors. There are no known sequence-specific binding motifs for mammalian PcG proteins; however, PcG proteins frequently bind to CpG islands (CGIs) [Oksuz et al., 2018]. Consistently, 95% (248/262) of PcG anchors overlap at least one CGI, and anchors frequently overlap clusters of nearby CGIs (Figs. S5, S6A). We mapped CGI clusters by merging CGIs within 5 kb, yielding a median cluster length of 4,827 bp for anchor-overlapping clusters versus a genome-wide CGI median of 585 bp (Fig. S6B).

Together, these results demonstrate that, in neurons, PRC1 preferentially occupies broad CGI-rich regions, likely facilitating the formation of long-range PcG contacts.

## 3 Discussion

We found that different subtypes of adult neurons exhibited distinct average intensities of PcG contacts. This pattern persisted even for contacts anchored at silent genes, indicating that differential expression of some PcG-anchor genes does not fully account for the observed differences and suggesting that other features of neuronal chromatin or nuclear organization likely contribute.

We found that PcG contacts anchor genes encoding transcription factors and neurotransmitter transporters characteristic of specific neuronal subtypes. Most of these genes, including *SOX2*, *DLX1/2*, and *SATB2*, formed contacts exclusively in either ENs or INs, while other genes, such as *FEZF2* and *MEIS2*, showed more intricate PcG-contact patterns.

We have demonstrated that PcG contacts emerged during development and reached peak intensity in adult neurons. Cortical neurogenesis is completed by the end of the second trimester; therefore, most of the time points we analyzed represented post-mitotic cells, suggesting that increasing PcG-contact intensity could reflect a gradual accumulation of Polycomb proteins on chromatin in post-mitotic neurons. Another contributing factor could be a redistribution of Polycomb from smaller to larger domains over time, an effect previously demonstrated in polymer simulations [Owen et al., 2023]. Such redistribution in long-lived post-mitotic cells may lead to Polycomb binding within a few very broad domains, giving rise to the bright PcG contacts observed in adult neurons.

Given the high affinity of PRC2 for CpG islands (CGIs), Polycomb is expected to accumulate at loci containing the longest CGIs, as we have shown for the adult cortex. Transcription factor (TF) genes are known to possess some of the longest CGIs [Elango and Yi, 2011]; therefore, such redistribution could help target Polycomb to TFs that must be repressed and maintained in a silent state.

The increase in PcG contact intensity coincided with reduced expression of transcription factors (TFs) at PcG anchors, indicating the functional importance of these contacts.

We did not detect unique PcG contacts in fetal or infant contact matrices, which may reflect either the lower average PcG signal or the smaller total number of contacts in the snm3C-seq matrices from Heffel et al. [Heffel et al., 2024].

An open question is the presence of ultra-long-range Polycomb-coupled interactions in other cell types and neuronal models. Interactions similar to what we observed in neurons were reported for mouse embryonic stem cells [Schoenfelder et al., 2015, Boyle et al., 2020] and human hematopoietic stem and progenitor cells [Zhang et al., 2020]. Interestingly, in iPSC-derived neurons and neuronal progenitor cells, the average frequency of PcG contacts is much lower than in neurons from post-mortem brain [Zagirova et al., 2025].

We and others have previously described the neuronal 3D genome as “domain-dominant”, characterized by pronounced topologically associating domains (TADs) and weak compartmentalization [Pletenev et al., 2024, Tian et al., 2023, Rahman et al., 2023, Heffel et al., 2024]. This observation is further supported by a recent single-cell atlas that found neuronal chromatin among the most domain-dominant across 16 human tissues [Zhou et al., 2025], raising the possibility that reduced compartmentalization facilitates PcG-contact formation or, conversely, that strong PcG interactions interfere with global compartmental organization.

## 4 Methods

### 4.1 RING1B ChIP-seq on human neuronal and non-neuronal nuclei

#### 4.1.1 Tissue collection and nuclei isolation

RING1B ChIP-seq was performed on neuronal and non-neuronal nuclei from postmortem human brain tissue in two biological replicates. The study was approved by the Bioethical Commission of the Institute of Gene Biology, Russian Academy of Sciences, and conducted in accordance with the Declaration of Helsinki. Human brain samples were provided by National BioService (St. Petersburg, Russia) with informed consent from donors or their next of kin; donor metadata are detailed in the Supplementary Materials (Table S2). Tissue blocks were excised from the Wernicke’s area (posterior Brodmann area 22, BA22p). Nuclei were extracted and sorted using fluorescence-activated nuclear sorting (FANS) as previously described in [Pletenev et al., 2024] with modifications.

In brief, approximately 600–800 mg of frozen tissue was sectioned into 200 mg pieces, collected using sterile tweezers, and immediately placed on dry ice. Tissue was then homogenized in 100 *µ*L of fixative solution (1× PBS, 1% formaldehyde, Sigma-Aldrich) using an automated pestle homogenizer. The homogenate was incubated for 10 min at room temperature in a total volume of 10 ml of fixative solution, and crosslinking was then quenched with glycine (final concentration 125 mM). Following centrifugation (1100 *g*, 5 min, 4 °C), the pellet was washed twice with NF1 buffer (10 mM Tris–HCl pH 8.0, 1 mM EDTA, 5 mM MgCl_2_, 100 mM sucrose, 0.5% Triton X-100) and incubated for 30 min in NF1 buffer. The suspension was homogenized on ice using Dounce homogenizer, and then filtered through a 70 *µ*m strainer. A sucrose cushion (1.6 M sucrose, 10 mM Tris-HCl pH 8.0, 3 mM MgCl_2_, 1 mM DTT) was layered under the homogenate, followed by centrifugation at 3200 *g* for 30 min at 4 °C with the brakes set to “low” to separate nuclei from debris.

Pelleted nuclei were sequentially washed with NF1 buffer and PBS, then resuspended in PBST containing 5% BSA and 3% fetal bovine serum. Nuclei were incubated overnight at 4 °C in the dark with a fluorescently conjugated NeuN antibody (Abcam, ab196184; 1:500, 100 *µ*L per million nuclei). The next day, nuclei were washed and resuspended in 1× PBS, and stained with DAPI (2 *µ*g/mL). After passing through a 35 *µ*m strainer, nuclei were sorted into NeuN-positive and NeuN-negative fractions using FANS. Sorted nuclei (approximately 1 million per sample) were collected in FANS buffer (1× PBS, 0.1% Tween-20, 5% BSA), centrifuged (2300 *g*, 10 min, 4 °C), washed with PBS, and either snap-frozen and stored at −80 °C or used immediately for ChIP-seq.

#### 4.1.2 Chromatin immunoprecipitation (ChIP)

The nuclei pellet was resuspended in 500 *µ*l RIPA buffer (50 mM Tris-HCl pH 8.0, 150 mM NaCl, 2 mM EDTA, 0.5% SDC (sodium deoxycholate), 0.3% SDS, 1% NP-40, 1× Protease Inhibitor Cocktail (the stock solution 100× Protease Inhibitor Cocktail (PIC), Bimake), 1×PMSF) and incubated for 15 min on ice. The cell lysate was sonicated on ice for 12 cycles: 30 sec pulse and 1.5 min off, using VirSonic 100, VirTis sonicator, to shear DNA to an average fragment size of 100 - 500 bp. After the sonication, the cell debris was pelleted for 5 min, 4 °C, 20000 *g*. The supernatant was transferred into Amicon 30K filter unit (Millipore), centrifuged for 5 min, 4 °C, 14000 *g*, and the chromatin was transferred into a new low-binding tube and diluted with 1 ml of ice-cold IP-buffer (0.5 M LiCl, 1% NP-40, 1% sodium deoxycholate, 100mM Tris-HCl pH 8.0, 1×PIC, 1×PMSF). 100 *µ*l (10% of the whole sample) were taken as input and frozen at-20 °C.

RING1B antibody (Active Motif, 39663; 1 *µ*g per million nuclei) was added, and samples were incubated overnight at 4 °C with rotation. Protein A/G magnetic beads (Thermo Scientific) were used to precipitate the chromatin-antibody complexes. Beads were washed with Beads Washing Buffer (1× RIPA, 0.5% BSA) as follows: twice for 15 min at 4 °C with 1 ml of buffer with rotation and then with 1 ml of buffer overnight at 4 °C. After washing, the beads were incubated with the chromatin-antibody mixture for 6 hours at 4 °C with rotation. The complexes were isolated on a magnetic stand as recommended (Life Technologies). The DNA was extracted with phenol:chloroform:isoamyl alcohol (25:24:1) and precipitated with 2.5 volumes of 96% EtOH (in the presence of 0.3M NaOAc and using glycogen and tRNA as co-precipitants). The pellets were dissolved in 50 *µ*l of 10 mM Tris-HCl (pH=8.0) and treated with RNase A (ThermoFisher Scientific, 10 *µ*g/*µ*l) for 30 minutes at 37 °C. After that, two volumes of AmpureXP magnetic beads (Beckman Coulter) were added to the samples. The mixture was incubated for 10 minutes at room temperature, followed by discarding the supernatant on a magnetic rack and washing the beads three times with 300 *µ*l of 80% ethanol. The beads were then resuspended in 50 *µ*l of 10 mM Tris-HCl (pH=8.0) and DNA was eluted at 52 °C for 15 minutes. After the incubation, the samples were placed on the magnetic rack again and the eluate was collected into fresh tubes. The obtained DNA in the amount of 10–100 ng was used for library preparation.

#### 4.1.3 ChIP-seq library preparation

End repair was performed in 100 *µ*L reactions by combining 50 *µ*L of ChIP DNA sample with reaction mix containing 25 *µ*L mQ water, 10 *µ*L 10× T4 DNA ligase buffer, 5 *µ*L 10 mM dNTP mix, 5 *µ*L T4 polynucleotide kinase (10 U/*µ*L, NEB), 4 *µ*L T4 DNA polymerase (3 U/*µ*L, NEB), and 1 *µ*L Klenow fragment (5 U/*µ*L, NEB). Reactions were incubated 30 min at room temperature and purified with 2 volumes of AMPure XP beads; DNA was eluted in 50 *µ*L 10 mM Tris-HCl.

For A-tailing, 30 *µ*L mQ water, 10 *µ*L 10× NEBuffer 2, 5 *µ*L 10 mM dATP, and 5 *µ*L Klenow fragment (exo*^−^*, 1 U/*µ*L, NEB) were added. Samples were incubated at 37 °C for 30 min and purified with 2 volumes of AMPure XP beads; DNA was eluted in 30 *µ*L 10 mM Tris-HCl.

Adapter ligation was performed by adding 1 *µ*L Illumina TruSeq adapters, 12 *µ*L mQ water, 5 *µ*L 10× T4 DNA ligase buffer (Thermo Scientific), and 2 *µ*L T4 DNA ligase (5 U/*µ*l, Thermo Scientific), followed by 16 h incubation at room temperature. Ligated DNA was purified with 1.2-1.5 volumes of AMPure XP beads and eluted in 30 *µ*L Tris-HCl.

PCR amplification of libraries was carried out with Illumina primers (forward: 5*^′^*-AATGATACGGCGACCACCGAGAT-3*^′^*; reverse: 5*^′^*-CAAGCAGAAGACGGCATACGA-3*^′^*) using KAPA HiFi HotStart polymerase. Cycling: 95 °C 3 min; 98 °C 20 s, 65 °C 15 s, 72 °C 20 s for the selected number of cycles; final extension at 72 °C for 3 min. The cycle number (typically 7–8 for input and 12–14 for ChIP) was chosen based on absence of overamplification on agarose gels and a DNA yield of ∼10–30 ng/*µ*l.

Libraries were sequenced on Illumina NovaSeq 6000 at the Skoltech Genomics Core, generating ∼40 million 100-bp paired-end reads per sample.

### 4.2 ChIP-seq data processing

#### 4.2.1 Adult excitatory and inhibitory cortical neurons

For automated annotation of PcG contacts, ChIP-seq data available under controlled access (samples H276 and H372) were used [Kozlenkov et al., 2018]. The mapping of reads and the subsequent identification of peaks were conducted utilizing nf-core/chipseq pipeline v2.0.0 [Patel et al., 2024a] of the nf-core collection of workflows [Ewels et al., 2020], utilizing reproducible software environments from the Bioconda [Grüning et al., 2018] and Biocontainers [da Veiga Leprevost et al., 2017] projects. The pipeline was executed with Nextflow v22.04.0 [Di Tommaso et al., 2017] with the following parameters: --genome hg38 --macs gsize 2700000000.

#### 4.2.2 Adult cortical neurons, oligodendrocytes, and NeuN(-) cells

Newly generated RING1B data together with H3K27me3 and BMI1 data available under controlled access [Loupe et al., 2024] were processed as following. Read mapping was conducted utilising nf-core/chipseq pipeline v2.1.0. The pipeline was executed with Nextflow v24.20.5 [Di Tommaso et al., 2017] with the following parameters: --macs gsize 2913022398 --fasta hg38.fa.gz-profile docker; fasta file was obtained from UCSC database. Final sets of peak coordinates were obtained using MACS3 v3.0.1 [Zhang et al., 2008] with BAM files for individual replicates as input files. To generate a merged bigWig file, BAM files for individual replicates were merged, sorted, and indexed using samtools v1.2 [Li et al., 2009]. Genome coverage was calculated from the merged BAM file using bedtools genomecov v2.30.0 [Quinlan and Hall, 2010], and the resulting bedGraph file was converted to the bigWig format using bedGraphToBigWig v455 [Kent et al., 2010]. Next, noise filtering with MACS3 bdgcmp function was applied to ChIP-seq data. The algorithm is based on comparing a bedGraph file with a control one using a Poisson model for detection and subtraction of the noise. The program was executed with the following parameters: callpeak --bdg and then bdgcmp-m ppois.

### 4.3 Bulk RNA-seq data processing

Bulk RNA-seq data available under controlled access for excitatory, inhibitory neurons, and oligodendrocytes were used [Kozlenkov et al., 2018]. Fastq files were processed using nf-core/rnaseq pipeline v3.14.0 [Patel et al., 2024b]. The pipeline was executed with Nextflow v23.10.1 with the following parameters: --genome hg38 --aligner star rsem --remove ribo rna.

Additionally, we used publicly available bulk RNA-seq data for NeuN(+) and NeuN(-) cells [Rizzardi et al., 2019]. Fastq files were processed using nf-core/rnaseq pipeline v3.8.1 [Patel et al., 2022] with Nextflow v22.04.0 with the following parameters: --genome hg38 --aligner hisat2

### 4.4 Snm3C-seq data processing

Contact pairs for cells of the developing and adult human cortex [Tian et al., 2023, Heffel et al., 2024] were downloaded from GEO database (GSE215353 and GSE213950). Cell-type annotations provided in the original studies were used. For the analysis, contact pairs were either aggregated from many cells into a pseudobulk contact matrix or converted into contact matrices for individual cells.

Pseudobulk matrices were generated using “cooler cload pairs” command of the cooler library v0.10.2 [Abdennur and Mirny, 2020]. Resulting matrices were normalized using iterative correction and eigenvector decomposition procedure [Imakaev et al., 2012] implemented in “cooler balance” command. For Tian et al. data, cells from all cortical regions were aggregated. For the majority of comparative analyses, matrices were sampled to the equal number of cells. Matrices were not sampled when analyzing multiple subtypes of adult neurons and when biological replicates were analyzed separately.

Contact matrices for the individual cells were created using “cooler cload pairs” command of the cooler library v0.10.2.

### 4.5 Manual annotation of PcG contact anchors

In this work, we refined and extended annotation obtained previously [Pletenev et al., 2024] by using snm3C-seq data with very high number of cells [Tian et al., 2023]. Manual annotation was performed using HiGlass visualization software v1.11.7 [Kerpedjiev et al., 2018] and two pseudobulk contact matrices with EN and IN cells respectively to better detect type-specific PcG contacts.

### 4.6 Automated annotation of PcG contacts

To extend manual annotation and detect PcG anchors that could have been missed, we used pseudobulk snm3C-seq contact matrices corresponding to cortical EN and IN at 100 kb resolution and applied logistic regression classifying pixels into “contact” and “not a contact” (S7A). To avoid many false positives and high computation time, we only considered pixels that overlap H3K27me3 ChIP-seq peak and at least one gene promoter with both their anchors.

#### 4.6.1 Selection of candidate regions

Candidate regions were identified from the H3K27me3 ChIP-seq dataset for adult EN and IN separately [Kozlenkov et al., 2018]. Peaks from two biological replicates were intersected using bedtools v2.30.0 [Quinlan and Hall, 2010]. The resulting consensus peaks were overlapped with gene promoters, defined as transcription start sites (GENCODE v41) extended by ±2*kb*. Peaks were then filtered to retain only those with a signal value ≥ 3.38 (approximately corresponding to the first quartile of the ChIP-seq peak intensity distribution) and peak length ≥ 1500 bp (S7B).

#### 4.6.2 Feature selection

Intrachromosomal pairwise interactions of candidate regions at genomic distance greater than 3 ∗ 10^6^ were selected for annotation. 21×21 windows centered on the candidate regions were collected from pseudobulk contact matrices using the Cooler library v0.9.3 [Abdennur and Mirny, 2020]. Windows with more than five empty rows or columns were filtered out. The following features were used in the model (S7C):

- central pixel;
- “central mean”: mean value of central 3*3 window;
- “central area n”: sum of central n*n window (n=3,5,9);
- “frame n”: mean value of a n*n perimeter of a square (n was varied from 5 to 21, including all odd numbers);
- “central to frame n”: central pixel divided by a frame n;
- “central to corners”: central pixel divided by mean value of four corner pixels.

#### 4.6.3 Logistic regression model

Logistic regression model from the scikit-learn library v1.5.2 was used [Pedregosa et al., 2011]. Statsmodels v0.14.4 was used for feature evaluation and selection [Seabold and Perktold, 2010]. Two models: for EN and IN data — were trained. Random 150 pairs from manually annotated anchors were selected and split into train (120) and test (30). To avoid overfitting, the train/test dataset was split by chromosomes, chromosomes 6, 14, and 15 were used for testing, while others were used for training. The following threshold values were used for classification: 0.65 for EN-based model and 0.8 for IN-based model.

#### 4.6.4 Combined manual and automated annotations

Although automated annotation outputs predicted PcG contacts, we did not use these predictions directly. Instead, we extracted anchors of predicted contacts and combined them with anchors from manual annotation. The assumption behind this operation is that PcG contacts are not specific, therefore if a locus forms contact with one Polycomb region, it must also form contacts with all the remaining Polycomb regions. The resulting anchors from manual and automated annotations have high overlap (S7D), and the union of these anchors was used as the final annotation (Table S1).

### 4.7 Aggregate peak analysis

Aggregate peak analysis was performed using coolpup.py library v1.1.0 [Flyamer et al., 2020]. 20 kb or 50 kb resolution was used. Mitochondrial and sex chromosomes were removed. When not stated otherwise, PcG contacts of anchors at genomic distances greater than 3 Mb were considered.

### 4.8 Aggregated analysis of PcG contacts

#### 4.8.1 Identification of PcG contacts in single cells

All intrachromosomal pairs of identified PcG anchors separated by more than 3 Mb were considered candidate PcG contacts in single cells. To remove low-signal candidates, we generated two pseudobulk contact matrices for cortical EN and IN. For each candidate, the “interaction intensity” and “background intensity” were computed as the total signal from the contact matrix over regions extending 10 kb and 50 kb from each anchor, respectively, divided by the corresponding area. Candidates with “interaction intensity” at least 1.5 times higher than “background intensity” in EN or IN matrix were retained for further analysis.

To identify PcG contacts in single-cell contact matrices, each “.tsv.gz” file with contact pairs obtained from [Tian et al., 2023] was converted into a “.cool” file representing contact matrix of a single cell using Cooler v0.10.2. PcG contacts intensities were extracted from contact matrices using the hictkpy v1.0.0 [Rossini and Paulsen, 2024].

#### 4.8.2 Aggregation and normalization

For each PcG contact, intensity was aggregated and normalized as follows:

- The “interaction intensity” was binarized (0 if no signal was detected, 1 otherwise).
- In each cell, the binary value was divided by the total number of contacts observed in that cell.
- For each cell type, the mean of these values was calculated across all cells of that type.
- The resulting values were then normalized to sum to one.

Cell types with fewer than 1500 cells, as well as non-cortical cell types, were excluded from the analysis.

#### 4.8.3 Visualization of aggregated PcG contacts

The aggregated and normalized intensities were clustered using the k-means algorithm implemented in scikit-learn v1.5.2 [Pedregosa et al., 2011]. Clusters were annotated manually. “Total” value for the heatmap was calculated as a mean value of PcG contacts in pseudobulk EN and IN contact matrices. Intensities of PcG contacts were also visualized using principal component analysis (PCA) from the scikit-learn.

### 4.9 Joint analysis of PcG contacts and gene expression in the adult brain dataset

To link Polycomb-mediated chromatin architecture with gene expression, aggregated PcG contacts were analyzed jointly with publicly available snRNA-seq data from the adult human brain [Siletti et al., 2023].

Gene expression data were analyzed using loompy v.3.0.7. Snm3C-seq and snRNAseq datasets were integrated manually based on cell-type annotations from the original studies (Table S3). Gene expression was log1p(CP10K)-normalized and averaged across cortical cell types. Gene activity was classified based on a manually selected threshold of 0.075: genes with mean expression above the threshold in all cell types were defined as “active”, those below it as “non-active”, and the remainder as “partially active”. For downstream analyses, we retained only active or partially active genes that were connected to non-active genes *via* PcG contacts.

For each gene, we calculated its average expression in EN and IN cell types and annotated it with the PcG contact clusters corresponding to the PcG contact it forms. Gene expression differences between EN and IN were assessed using the Wilcoxon signed-rank test implemented in the pingouin package (v0.5.5) [Vallat, 2018].

For each gene, expression and the intensities of the corresponding PcG contacts were calculated within each cell type. The Pearson correlation coefficient was then computed between mean gene expression and mean PcG contact intensity across cell types.

We hypothesized that for genes regulated by Polycomb, either gene expression or PcG contact should be observed. We aimed to identify two biologically meaningful cases of the “expression-or-loop” pattern: (1) those associated with differences between excitatory and inhibitory neurons, and (2) those explained by specific features of a single neuronal subtype. To distinguish between these cases, we designed two metrics. The first metric, based on mean gene expression across subtypes, was computed as the difference between the highest and second-highest expression values. For the second metric, we applied binary k-means clustering (k = 2) using gene expression and loop intensity values, and then calculated the Pearson correlation coefficient between predefined cell-type labels (EN/IN) and unsupervised cluster labels (from k-means). By evaluating these two metrics (S8), we manually selected genes that followed either form of the “expression-or-loop” pattern.

#### 4.9.1 Analysis of snRNA-seq data for the developing brain

To analyze gene expression in the developing human brain, we utilized publicly available snRNA-seq data [Herring et al., 2022]. Processed expression counts (file RNA-all full-counts-and-downsampled-CPM.h5ad) were downloaded from https://brain.listerlab.org and analyzed with Scanpy v1.10.1 [Wolf et al., 2018]. Samples were divided into four groups (Table S4), and counts were aggregated into four pseudobulk samples followed by log1pCPM normalization. For cell-type-specific analysis, counts were aggregated into pseudobulk EN and IN samples based on annotation from the original study.

Differential expression analysis was conducted using PyDESeq2 v0.5.0 [Muzellec et al., 2023] with default parameters, except for setting independent filter=False. 2T and 3T samples were grouped into a single “fetal” category due to low number of samples.

### 4.10 ChIP-seq data analysis

For analysis of ChIP-seq signal at gene promoters shown at Figures 4 and S5, GENCODE annotation v41 [Frankish et al., 2021] was used. Gene transcription start sites (TSSs) were extended by 20 kb to each side and overlapped with ChIP-seq peaks from six experiments; regions that overlapped at least one peak were kept. Bins overlapping either a 50-kb window around low-signal regions in EN/IN contact maps or regions listed in the ENCODE blacklist were removed. For the remaining regions, the ChIP-seq signal (-log10(Poisson p-value)) was summed over each region for each factor and Z-score normalized for k-means clustering (k=10). A heatmap was constructed by dividing each region into 100 bins; regions corresponding to genes on the reversed strand were mirrored. Clusters were categorized as”Polycomb-associated” if they had high fraction of regions overlapping PcG anchors. The average ChIP-seq signal across all regions was calculated for each of the 100 bins to visualize peak intensity and width. Differences in signal intensity at central bins between clusters were assessed using a two-sided Wilcoxon rank-sum test.

For annotation-independent analysis of PcG contacts, we calculated the average ChIP-seq signal for H3K27me3, BMI1, and RING1B in neurons in 10-kb genomic bins using deeptools multiBigwigSummary v3.5.5 [Ramírez et al., 2014]. Bins overlapping 50kb-neighborhood of regions of low signal in EN/IN contact maps or the ENCODE blacklist were excluded. The bins were then sorted by their average signal in ascending order. Inflection points, beyond which the signal values increased sharply, were identified on the resulting plots and bins were classified into “high signal” and “low signal”. High-signal bins were divided into 20 quantiles, and the fraction overlapping long-distance PcG anchors was determined for each quantile. Next, for each quantile, we selected corresponding bins and computed average contact frequency matrices for all intra-chromosomal bin pairs located 3 Mb apart.

### 4.11 CTCF-mediated loop annotation

CTCF ChIP-seq peak coordinates (DLPFC) from the original study [Loupe et al., 2024] were used. CTCF peaks overlapping either a 50-kb window around low-signal regions in EN/IN contact maps or regions listed in the ENCODE blacklist were excluded. Chromatin loops were annotated on contact maps from EN and IN at 2 kb and 5 kb resolutions using Chromosight v1.6.3 with default parameters [Matthey-Doret et al., 2020]. The resulting loops overlapping with CTCF peaks were considered CTCF-mediated loops.

### 4.12 Fetal H3K27me3 ChIP-seq data analysis

Publicly available chromHMM annotation based on ChIP-seq data for histone modifications in fetal brain was used [Rahman et al., 2023].

## Acknowledgements

We are grateful to the Core Centrum of IDB RAS and Skoltech BioImaging and Spectroscopy Core Facility for the excellent technical assistance. We thank the Center for Precision Genome Editing and Genetic Technologies for Biomedicine, IGB RAS for the cell sorting. We thank Skoltech Genomics Core Facility and Dr. Elena Shagimardanova for sequencing and quality control of ChIP-seq libraries. We thank National BioService Russian Biospecimen CRO (St. Petersburg, Russia) for providing brain samples. Some of the ChIP–seq and RNA–seq data used in this study were obtained from the NIMH Repository & Genomics Resource, a centralized national biorepository for genetic studies of psychiatric disorders.

## Declarations

### Data availability

ChIP-seq data generated in this study are available in the GEO database under the accession number GSE309367.

### Code availability

Code generated for this study is available at https://github.com/i-pletenev/polycomb_in_single_neurons_paper

### Author contribution

Ilya A. Pletenev: Formal analysis, Writing — original draft, Writing — review & editing. Nikita Vaulin: Formal analysis, Writing — original draft, Writing — review & editing. Maria N. Molodova: Investigation, Writing — review & editing. Ekaterina Kuznechenkova: Formal analysis, Writing — review & editing. Anastasia Soldatenkova: Formal analysis, Writing — review & editing. Olga I. Efimova: Investigation, Writing — review & editing. Anna V. Tvorogova: Investigation, Writing — review & editing. Philipp Khaitovich: Resources, Writing — review & editing. Sergey V. Razin: Resources, Writing — review & editing. Sergey V. Ulianov: Supervision, Writing — original draft, Writing — review & editing. Ekaterina E. Khrameeva: Conceptualization, Funding Acquisition, Supervision, Writing — original draft, Writing — review & editing.

