## Supplemental Figures and Tables 2-4 for "Ultra-long-range Polycomb-coupled interactions underlie subtype identity of human cortical neurons"

### Supplementary materials

**Table S2** RING1B ChIP-seq sample metadata.

| Cell Type | Donor | Sex | Age, years | Total number of reads (antibody) | Total number of reads (input control) |
|---|---|---|---|---|---|
| NeuN+ | H7 | female | 28 | 72,437,865 | 71,492,746 |
| NeuN+ | H8 | male | 67 | 60,550,970 | 58,955,823 |
| NeuN- | H56 | female | 46 | 60,616,638 | 93,234,637 |
| NeuN- | H8 | male | 67 | 54,013,971 | 55,243,503 |

**Table S3** Manual integration of snm3C-seq cell-type annotation from Tian et al. [Tian et al., 2023] with scRNA-seq cell-type annotation from Siletti et al. [Siletti et al., 2023].

| **Tian_MajorType** | **Siletti_Cluster_ID** |
|---|---|
| Amy-Exc | 153, 154, 155, 156, 157, 158, 159, 160, 161, 162 |
| L2/3-IT | 120, 121, 122, 123, 124, 125, 126, 127, 128, 129, 130, 131, 134 |
| L4-IT | 138, 141 |
| L5-ET | - |
| L5-IT | 142, 143, 144 |
| L5/6-NP | 86, 87, 88, 89, 90, 91, 92, 93, 94, 95, 96 |
| L6-CT | 109, 110 |
| L6-IT | 148, 149 |
| L6-IT-Car3 | 150, 151 |
| L6-IT-Car3 | 152 |
| L6b | 98, 99, 101, 103, 105, 106 |
| Lamp5 | 286, 287, 288 |
| Lamp5-Lhx6 | 270, 271, 272, 274, 275 |
| Pvalb | 240, 255, 256, 257, 258, 259, 260, 261, 262, 263 |
| Pvalb-ChC | 264, 265 |
| Sncg | 282, 283 |
| Sst | 241, 242, 243, 248, 249, 250, 251, 252, 253, 254 |
| Vip | 284, 285, 289, 290, 291, 292, 293, 294, 295, 296 |
| Chd7 | - |
| Foxp2 | 229 |
| MSN-D1 | 209, 210, 211, 214, 215, 219, 220, 227, 228, 230, 231, 232, 234 |
| MSN-D2 | 207, 208, 212, 213, 216, 217, 221 |
| SubCtx-Cplx | - |
| ASC | 52, 53, 54, 55, 56, 57, 58, 59, 60, 61, 62, 63, 64 |
| OPC | 32, 33, 34, 35, 36 |
| ODC | 40, 44, 45, 46, 47, 48, 49, 50 |
| MGC | 4, 5, 6, 7, 8, 9, 10, 11, 12 |
| PC | - |
| VLMC | 21, 22, 23, 25, 26, 27, 28 |
| EC | 14, 15, 16, 17 |

**Table S4** Correspondence between Herring et al. sample age and snm3C-seq age group.

| **Sample age** | **Age group** |
|---|---|
| ga22 | 2T |
| ga24 | 2T |
| ga34 | 3T |
| 118d | Infant |
| 179d | Infant |
| 20yr | Adult |
| 25yr | Adult |

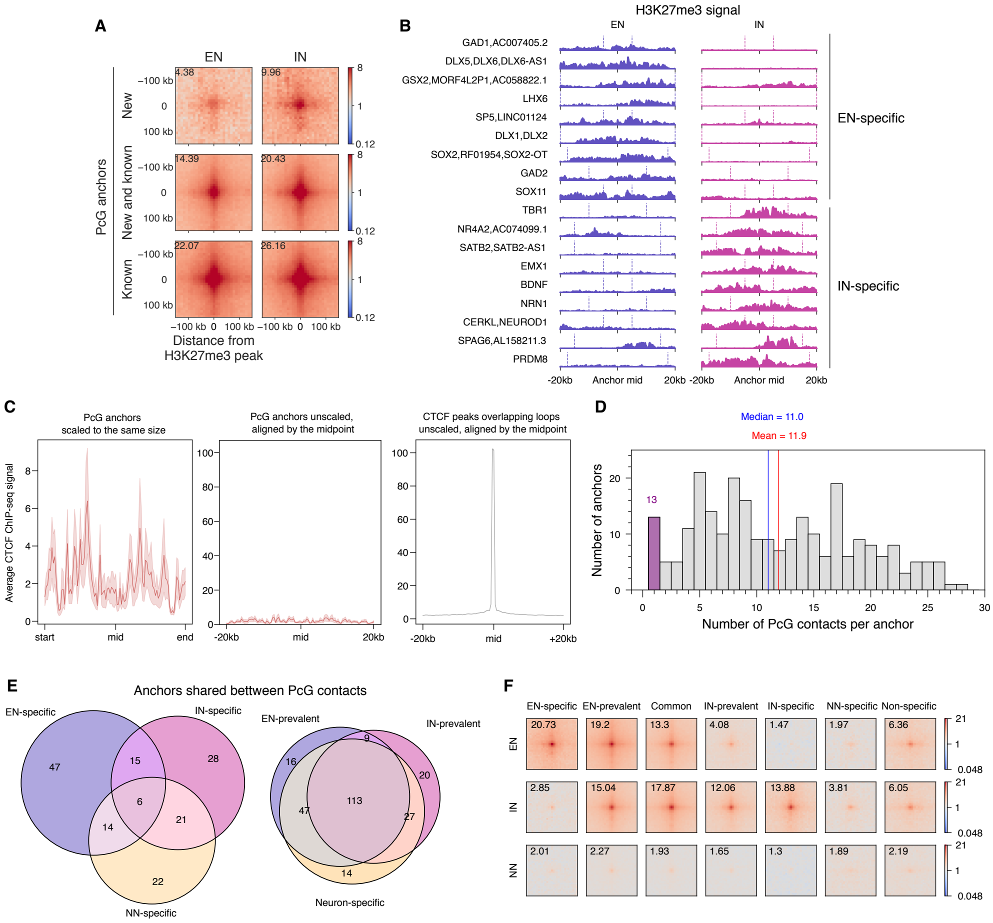


**Fig. S1** A. Average PcG contacts detected by automated annotation on EN and IN contact matrices. Contacts are divided into groups based on whether both contact anchors were present in manual annotation ("Known"), not present ("new"), or only one of two anchors was present ("New and known"). Values are normalized by expected contact frequency at given distance. B. H3K27me3 ChIP-seq signal centered at the midpoints of PcG anchors forming EN-specific and IN-specific PcG contacts. Vertical lines represent the actual boundaries of a PcG anchor. The boundaries of DLX5/6 locus extend beyond shown region. Genes overlapping anchors are depicted on the left. Vertical axis scale is equal for all tracks. C. Comparison of PcG anchors and CTCF-mediated loops. Average CTCF ChIP-seq signal was calculated for PcG anchors scaled to the same size (left), unscaled PcG anchors aligned by the midpoint (center), and CTCF peaks anchoring CTCF loops (right). Area around the line — standard error of the mean. D. Distribution of the number of PcG contacts per anchor. E. PcG anchors grouped by the clusters that corresponding contacts belong to. For example, if a region anchors one contact from EN-specific cluster and another — from IN-specific, it will be counted at the intersection of these two sets. F. Average PcG contact frequency per cluster calculated from pseudobulk snm3C-seq data and normalized by expected signal at a given genomic distance. Same as in Figure 1C, but including NN-specific and Non-specific clusters.

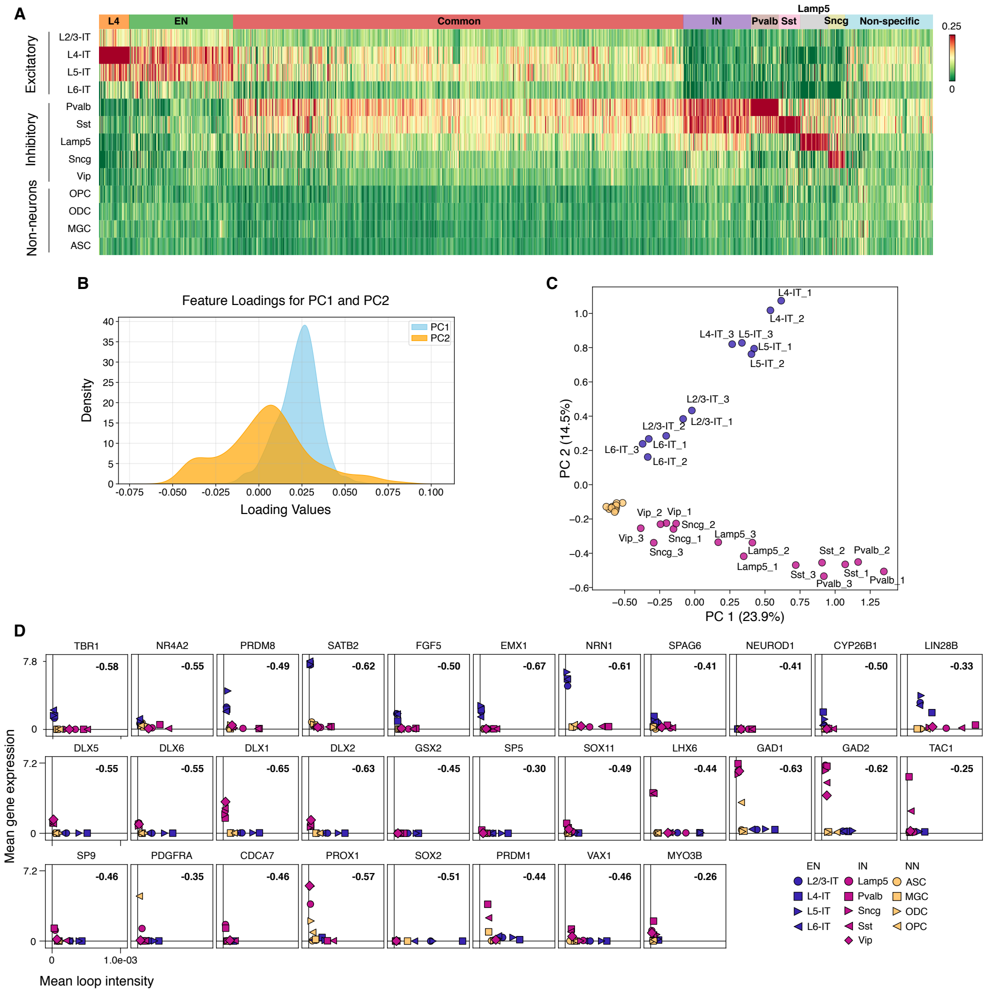


**Fig. S2** A. Normalized number of cells containing a PcG contact in the brain cell types. Each column corresponds to a separate PcG contact. B. PCA loadings corresponding to Figure 2A. C. PCA based on PcG contact intensity calculated for neuronal subtypes and non-neurons; for three biological replicates separately. D Mean PcG contact intensity vs mean gene expression of a gene overlapping PcG anchor; for genes from Figure 2C. Pearson correlation coefficient (r) is shown in the corner. “ASC” — astrocytes, “MGC” — microglia, “ODC” — oligodendrocytes, “OPC” — oligodendrocyte progentor cells.

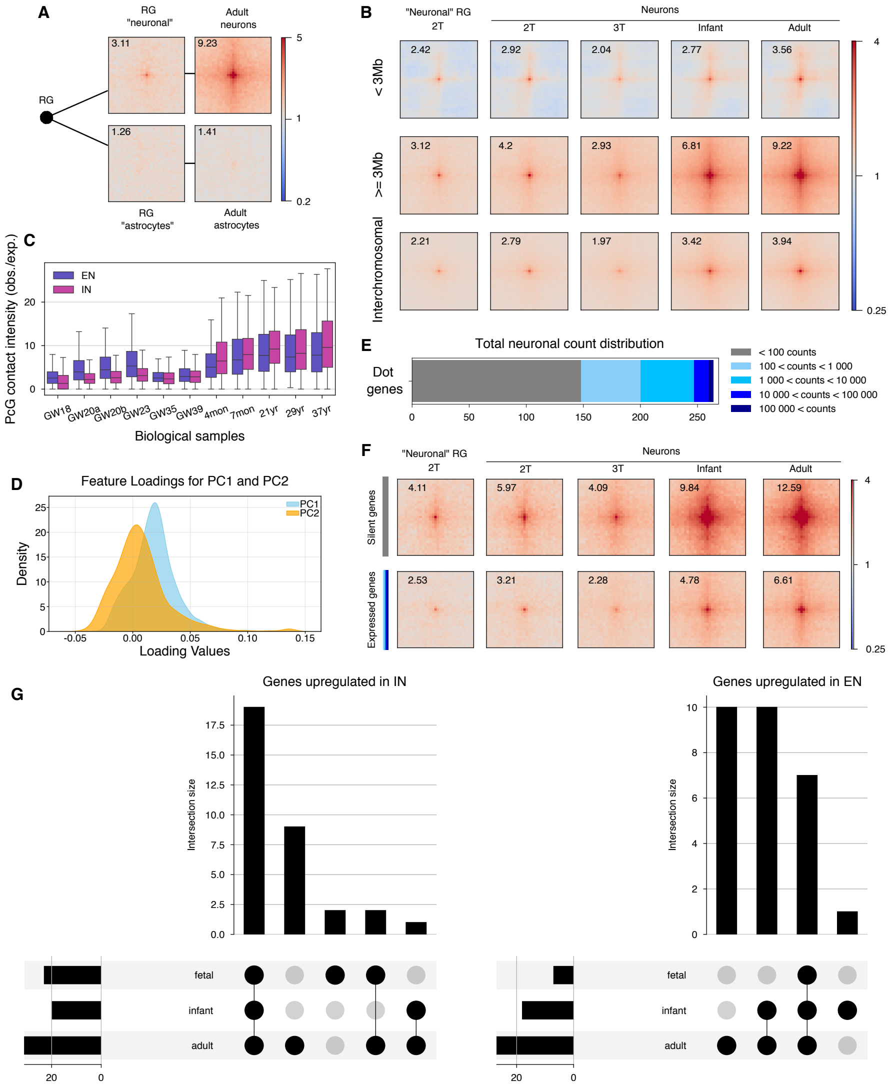


**Fig. S3** A. The average snm3C-seq (pseudobulk) signal (observed over expected at a given genomic distance) of PcG contacts for two clusters of radial glial cells (RG) compared with adult neurons and astrocytes. Annotation of RG clusters and cell types from original study was used [Heffel et al., 2024]. B. The average snm3C-seq (pseudobulk) signal (observed over expected) of PcG contacts for "neuronal" cluster of RG and neurons at four time points. Contacts are grouped by genomic distance between anchors. C. Distribution of PcG contact intensity for separate biological samples. "GW" — gestation week, "mon" — month, "yr" — year. D. PCA loadings for PC1 and PC2 from S3C. E. Total number of expression counts summarized by four time points for PcG anchor genes. "¡ 100 counts" (silent genes) is a set of genes that either were filtered out in the original study [Herring et al., 2022] due to low count number, or had less than 100 counts. F. The average snm3C-seq (pseudobulk) signal (observed over expected) of PcG contacts for "neuronal" cluster of RG and neurons at four time points. PcG contacts are formed by anchors containing genes silent at all stages (top) or expressed at least in one stage (bottom). G. Upset plot for genes differentially expressed between EN and IN at any developmental stage. Categories represent stage at which a gene is differentially expressed. Stages 2T and 3T were merged into "fetal" due to low sample number.

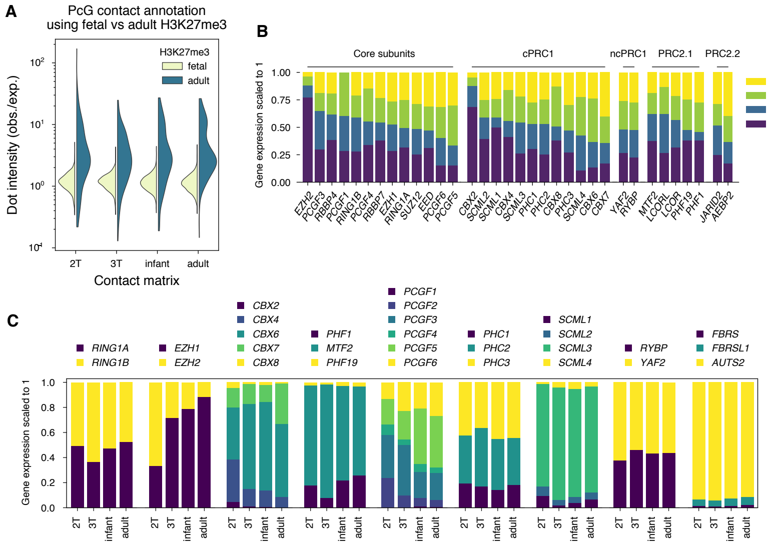


**Fig. S4** A. Intensity of PcG contacts annotated in adult neuronal data compared with annotation using all pairs of fetal-specific H3K27me3-occupied regions. Intensity is normalized by the expected contact frequency at a given genomic distance. B. Relative expression of Polycomb genes at developmental stages. C. Relative expression of Polycomb genes separated into groups of paralogs.

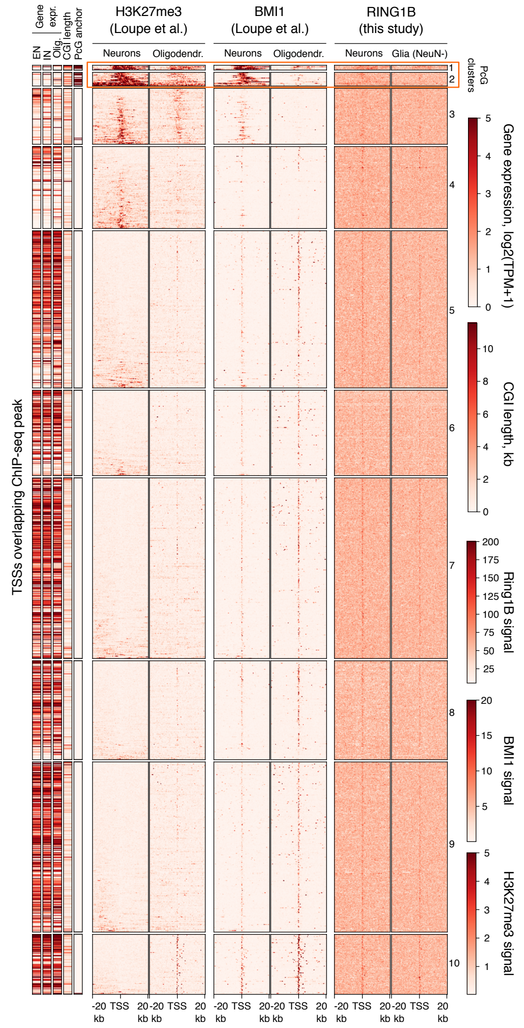


**Fig. S5** ChIP-seq signal at broad loci around gene TSSs. Only TSSs that overlap ChIP-seq peak in at least one experiment are considered. Additional annotation include: an overlap between TSS and the annotation of PcG contact anchors, the length of CGI in the region near TSS (+-5kb), and gene expression in adult cortical EN, IN, and oligodendrocytes [Kozlenkov et al., 2018].

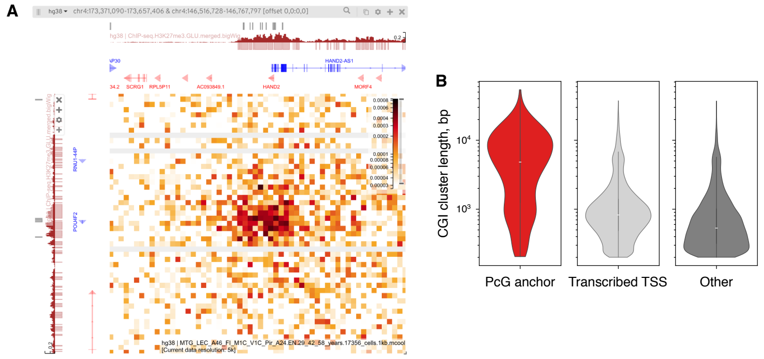


**Fig. S6** A. Genome browser view depicting a PcG contact between *HAND2* and *POU4F2* loci for excitatory neurons. Red tracks to the left and top — H3K27me3 data. Grey rectangles — CGI coordinates. B. CGI cluster length for CGIs overlapping PcG anchors, transcribed TSSs, or none of the above. CGIs intersecting poorly mappable genomic regions were excluded.

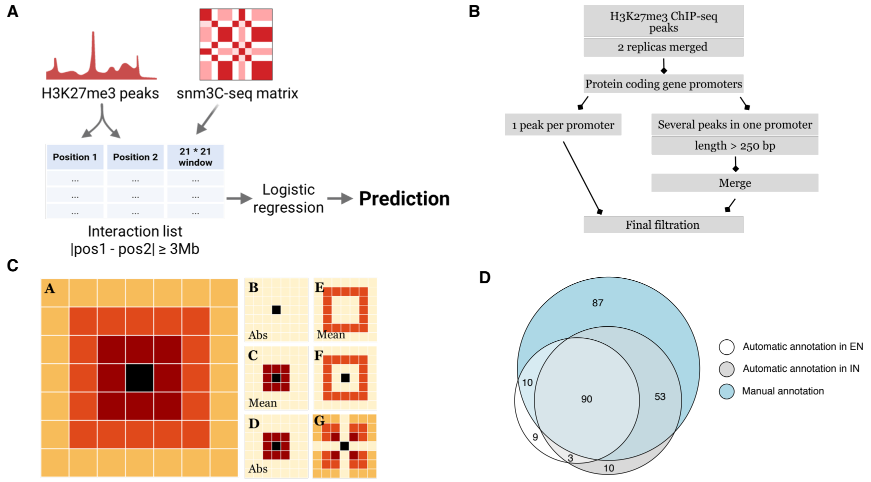


**Fig. S7** A. An overview of automated annotation of PcG contacts. B. An algorithm of H3K27me3 ChIP-seq peak filtration used for automated annotation. C. Features used for automated annotation. D. Average PcG contacts detected by automated annotation on EN and IN contact matrices. Contacts are divided into groups based on whether the contact anchors were present in manual annotation ("known") or not ("new"). Values are normalized by expected contact frequency at given distance.

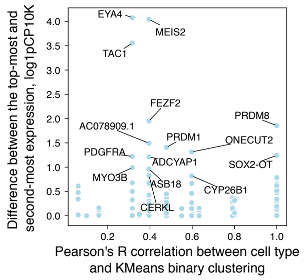


**Fig. S8** Two metrics of "expression-or-loop" pattern: difference between the top-most and second-most expression (across subtypes) and Pearson's correlation between cell subtype and KMeans binary clustering.
